# First Record of the Ranellid Gastropod Turritriton labiosus (Wood, 1828) (Gastropoda: Cymatiidae) from Tenerife, Canary Islands (NE Atlantic)

**DOI:** 10.1101/2025.07.08.663791

**Authors:** Michael Bommerer

## Abstract

We report the first confirmed record of the ranellid gastropod Turritriton labiosus (Wood, 1828) from the island of Tenerife in the northeastern Atlantic Ocean. A single shell specimen was discovered at 6 m depth approximately 100m off the southern coast. The specimen matches the reddish-orange morphotype previously reported from Indo-Pacific localities such as Guam and the Marshall Islands. This constitutes a new biogeographic record, potentially indicative of long-distance larval dispersal or an unrecognized Atlantic population. Morphological comparisons, shell dimensions, and habitat context are discussed. The specimen is housed in the author’s curated private scientific reference collection and has been documented on iNaturalist and RedPROMAR.

## Introduction

Turritriton labiosus (Wood, 1828) is a predatory marine gastropod belonging to the family Cymatiidae. The species is widely distributed throughout the Indo-Pacific, ranging from the Red Sea and western Indian Ocean to the central Pacific, including Japan, Micronesia, and Hawaii, and is typically associated with coral reef and sandy substrates in tropical and subtropical waters. In the eastern Atlantic, its presence has historically been regarded as doubtful or unconfirmed due to the lack of verifiable specimens.

A shell specimen from Lanzarote was first documented by Vicente (2007), representing the earliest modern confirmed record of T. labiosus in the Canary Islands. Unverified mentions on citizen science platforms and regional faunal databases—such as the Biota database (Biodiversidad de Canarias, 2025)—have suggested a possible occurrence in Macaronesia. However, the absence of voucher material (i.e., preserved physical specimens deposited in curated collections) has thus far precluded formal confirmation in biodiversity systems such as iNaturalist and RedPROMAR.

Historical references to Turritriton labiosus in the eastern Atlantic, including the Canary and Cape Verde Islands, appeared in the malacological literature of the late 20th century (Nordsieck & García-Talavera, 1979; Cosel, 1982; García-Talavera, 1983, 1987). While these records are of historical interest, they lack verifiable voucher specimens, diagnostic imagery, or precise locality data. As such, they have not been incorporated into modern biodiversity platforms, including iNaturalist, RedPROMAR, or WoRMS. A single user-submitted image of an eroded shell from Tenerife exists on WoRMS (Kapella, 2023), but it remains taxonomically unverified and is not included in the platform’s curated distribution data.

In addition to confirmed occurrences in the eastern Atlantic, a number of records tentatively identified as Turritriton labiosus have been reported from the western Atlantic, including coastal regions of Florida, Cuba, and St. Kitts & Nevis. These records, derived primarily from georeferenced observations submitted to citizen science platforms such as iNaturalist (iNaturalist, 2025), lack associated voucher specimens and have not undergone formal morphological or molecular validation. Preliminary comparisons suggest that western Atlantic shells may differ in overall profile and sculpture from both the Tenerife specimen and Indo-Pacific morphotypes. This raises the possibility that these observations represent a distinct taxon rather than part of a continuous trans-Atlantic distribution. Further anatomical and genetic analyses will be required to determine whether they represent misidentified T. labiosus, regionally diverging populations, or an undescribed western Atlantic species.

We present the first confirmed specimen of T. labiosus from Tenerife, Canary Islands, expanding the known distribution of the species within the Canary Archipelago. We provide a detailed morphological analysis of the specimen, compare it with Indo-Pacific morphotypes, and briefly evaluate its implications for understanding the species’ Atlantic distribution.

## Material and Methods

On 24 May 2025, a single specimen of Turritriton labiosus (Wood, 1828) was collected during a shallow SCUBA survey off the southern coast of Tenerife, Canary Islands (28°01’40”N, 16°34’06”W), at a depth of approximately 6 meters. The specimen was observed aperture-up on a sandy substrate plateau adjacent to a rocky reef system. The shell was inhabited by a hermit crab, identified in situ as Pagurus anachoretus (Risso, 1816), a species commonly recorded in the eastern Atlantic and Mediterranean Sea (WoRMS, 2024).

The hermit crab was carefully extracted underwater, after which the shell was gently cleaned using seawater and a soft-bristled brush. Residual organic material was removed by immersion in a diluted sodium hypochlorite solution (1 part sodium hypochlorite to 4 part water) for one hour. All morphological measurements were taken using calipers to the nearest 0.1 mm.

Comparative analysis was conducted using high-resolution images and documented specimens of Indo-Pacific T. labiosus from published literature and curated online reference databases. The Tenerife specimen is housed in the author’s private scientific reference collection (Tenerife, Canary Islands), maintained under professional curation standards in accordance with accepted malacological practices. Photographic documentation has been archived on iNaturalist and RedPROMAR to support biodiversity monitoring and regional faunal database

## Taxonomy

### Phylum

***Mollusca****Cuvier, 1797*

### Class

***Gastropoda****Cuvier, 1795*

### Subclass

***Caenogastropoda****Cox, 1960*

### Order

***Littorinimorpha****Golikov & Starobogatov, 1975 (sensu traditional classification; see Bouchet et al. 2017 for alternative rank-free placement)*

### Superfamily

***Tonnoidea****Suter, 1913*

### Family

***Cymatiidae****Iredale, 1913*

### Genus

***Turritriton****Dall, 1904*

### Species

***Turritriton labiosus****(Wood, 1828) (Fig. 1–5)*

## Results

### Shell morphology

The shell is solid, thick, and broadly fusiform, measuring 23.3 mm in total length and 16.8 mm in maximum width. The body whorl accounts for more than 50% of the total shell length, with a moderately extended, tapering siphonal canal.

**Figure 1.**
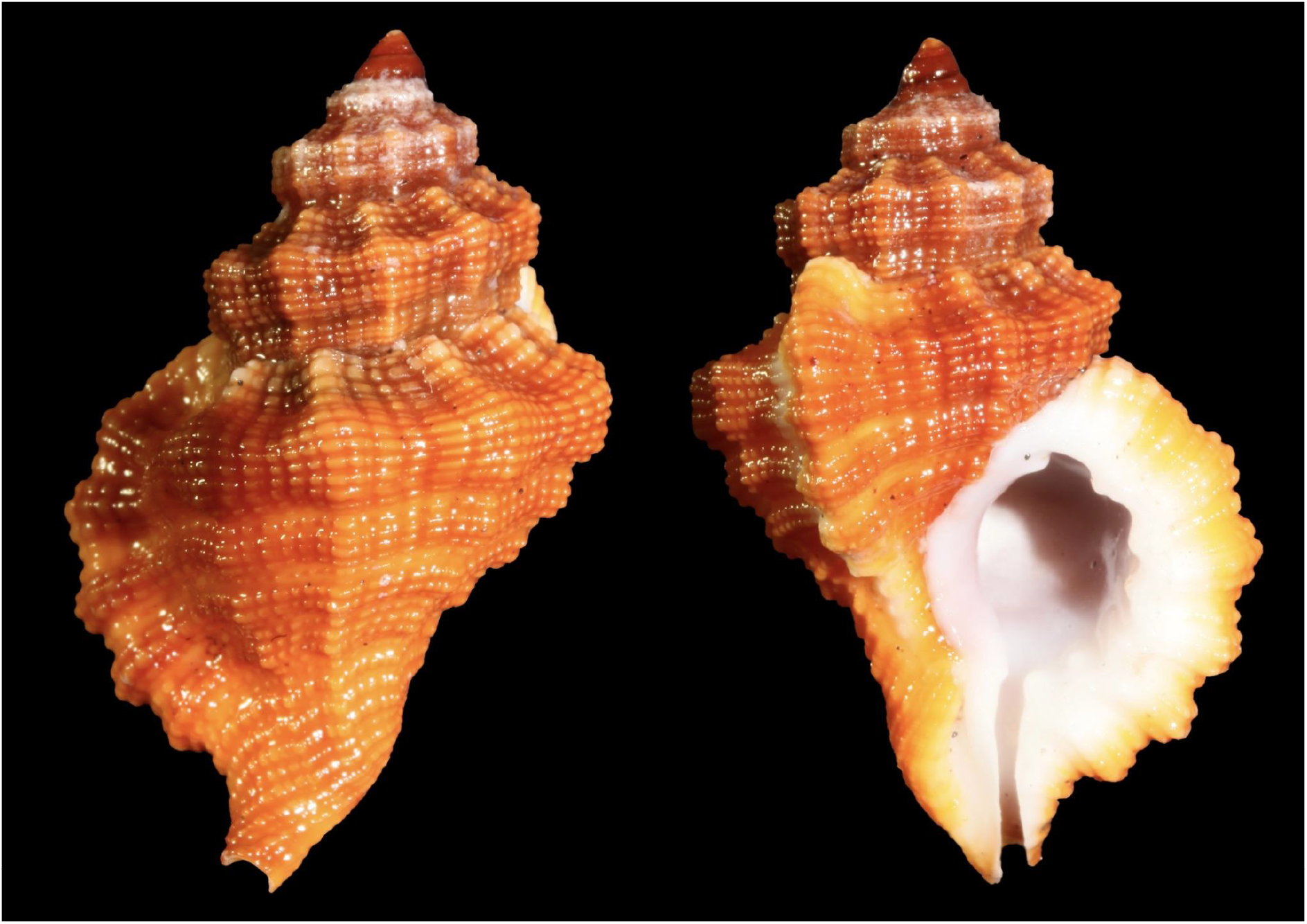
*Turritriton labiosus* (Wood, 1828) from Tenerife, Canary Islands. Shell collected at 6 m depth on 24 May 2025, 140m offshore southern Tenerife (28°01′40″N, 16°34′06″W). Shell length: 23.3 mm; width: 16.8 mm. The specimen displays a vivid reddish-orange ground color, prominent cancellate sculpture formed by intersecting axial ribs and spiral cords, and a faint whitish subsutural band present on each whorl.

### Protoconch

The protoconch is intact and comprises 2–3 smooth, rounded whorls. Its morphology is consistent with the planktotrophic larval type reported for Turritriton labiosus and related taxa.

### Teleoconch

The teleoconch consists of approximately four convex whorls, ornamented with strongly cancellate sculpture formed by rounded axial ribs intersected by raised spiral cords. Nodules are prominent at the intersections, particularly near the shoulder, giving a distinctly tuberculate appearance to the whorl profile.

### Aperture

The aperture is broad and ovate. The outer lip is moderately thickened and slightly flared, bearing six strong, evenly spaced white labial denticles on the inner edge of the outer lip. The columella is smooth, white, and gently twisted. The siphonal canal is moderately long, open, and curved slightly to the left. No operculum was preserved with the specimen.

### Periostracum

A thin, translucent, and tightly adherent periostracum covers most of the shell surface. No setae, lamellae, or additional surface ornamentation were observed.

### Coloration

The shell exhibits a uniform reddish-orange to reddish-brown ground color, extending over both the spire and body whorl. The aperture, columella, and siphonal canal are glossy white. A faint, irregular whitish subsutural band is present on each whorl, most prominently near the apex. This feature closely resembles the pale sutural banding reported in dark-pigmented morphs from Kwajalein Atoll (Johnson, 2020).

## Conclusion and Discussion

The specimen of Turritriton labiosus collected from Tenerife represents the first verified occurrence of this species on the island and constitutes the second confirmed record for the Canary Islands and the northeastern Atlantic Ocean (Fig. 1). Previously known from Indo-Pacific localities including Micronesia, Guam, and the Marshall Islands, T. labiosus is characterized by its robust axial sculpture, low-spired profile, and vivid reddish-orange coloration—features consistent with morphotypes described from Apra Harbor, Guam (Abela, 2018; Fig. 2), and “Sp. 2” from Kwajalein Atoll (Johnson, 2020; Figs. 3–5).

**Figure 2.**
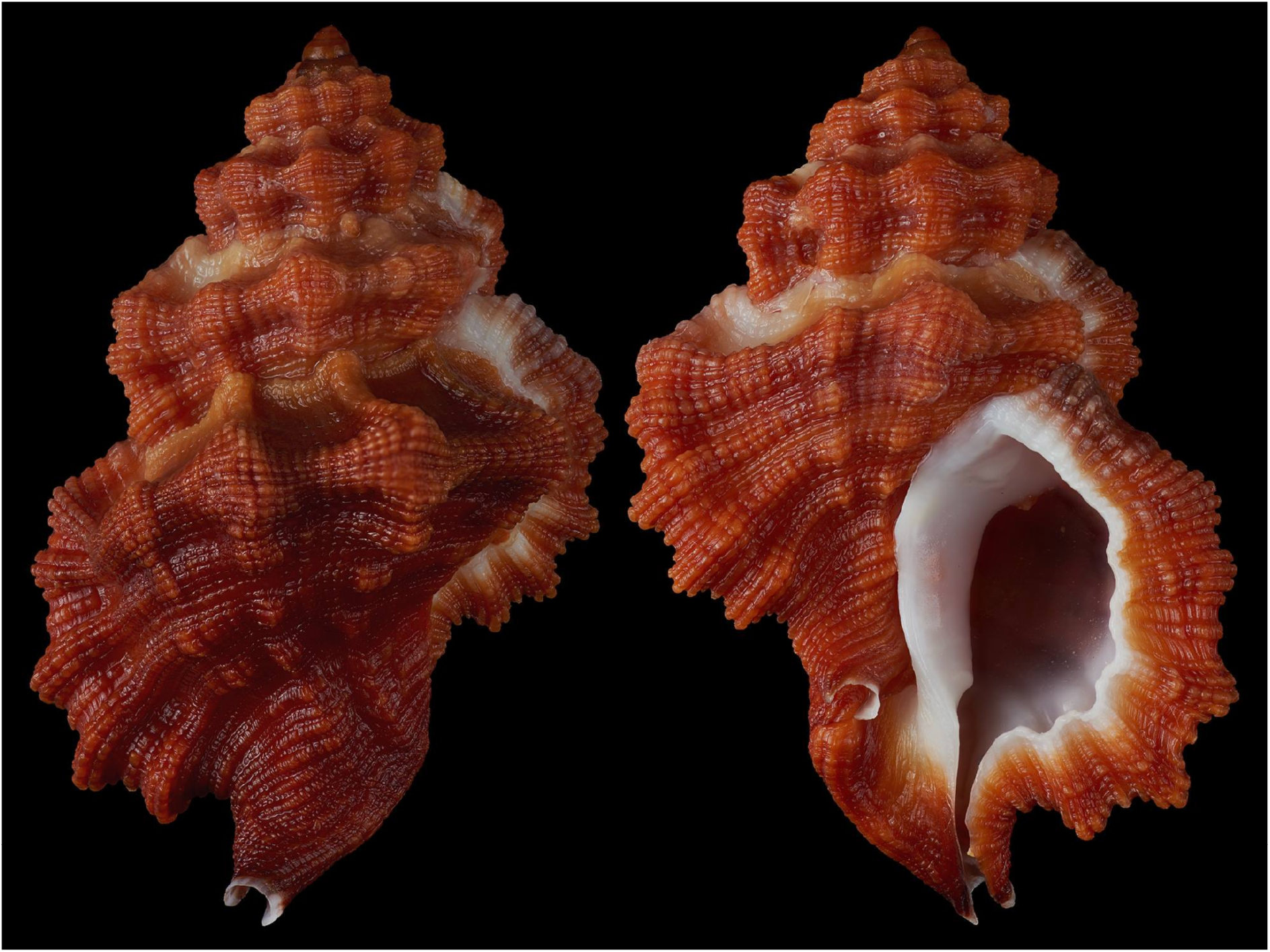
*Turritriton labiosus* (W. Wood, 1828), Apra Harbor, Guam. Shell (28.9 × 19.0 mm) collected dead at ∼45 ft on silty sand, 20 October 2018. This reddish-orange morph exhibits strong morphological congruence with the Tenerife specimen, particularly in axial sculpture and overall shell pigmentation. Image © Bob Abela, reproduced from www.bobabela.com, (used with permission)

**Figure 3.**
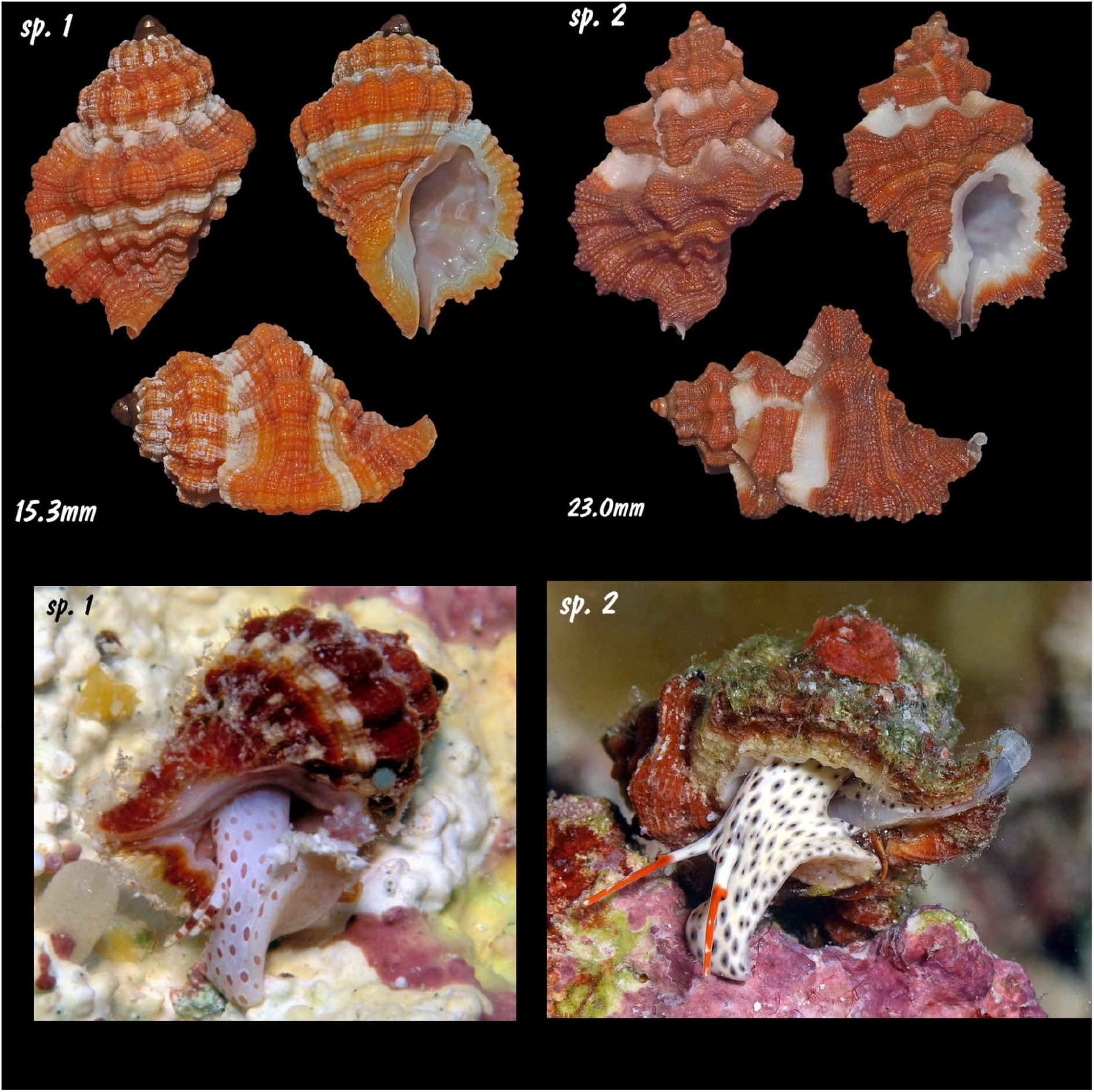
Comparison of two morphotypes previously grouped under *Turritriton labiosus*, photographed alive and post-collected by Scott Johnson at Kwajalein Atoll, Marshall Islands. Top: Shell morphology of “sp. 1” (15.3 mm) and “sp. 2” (23.0 mm), showing finer axial ribs in sp. 1 and coarser, widely spaced ribs in sp. 2. Bottom: Live views of the same specimens, highlighting differences in soft-part pigmentation. Sp. 1 exhibits a pale bluish-violet foot with diffuse purplish-red maculations and longitudinally striped cephalic tentacles of deep red to purple hues. Sp. 2 displays a nearly white foot uniformly covered with distinct black maculae, and strikingly colored cephalic tentacles with vivid orange distal tips. These soft-part patterns are consistent across observed individuals and support Johnson’s (n.d.) hypothesis of significant interspecific variation. Johnson (Facebook post, n.d.) observed: “The animals are quite different… I’ve not seen that much animal color variation in any of the tritons,” and raised the possibility that these may represent distinct species rather than intraspecific variation within T. labiosus.

**Figure 4.**
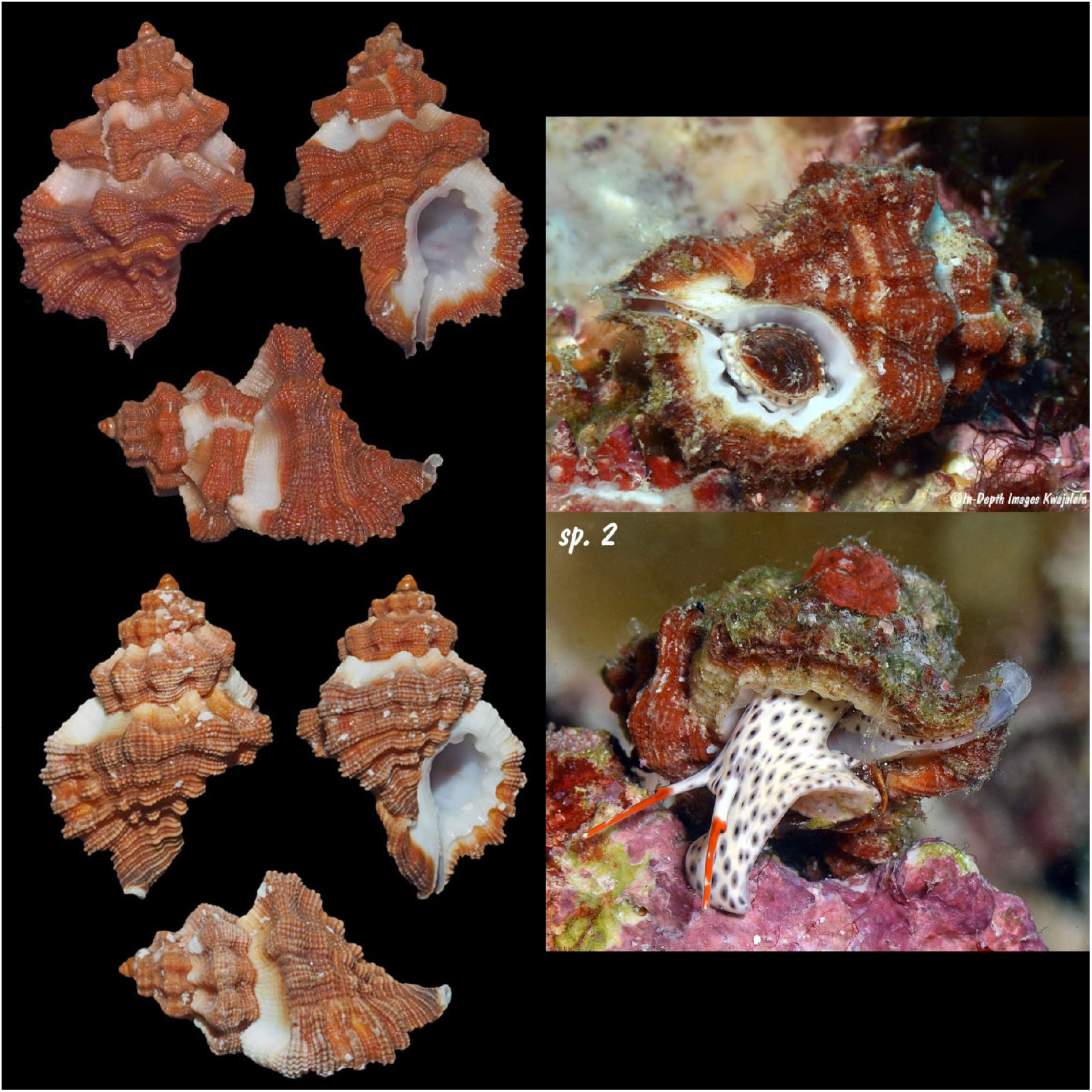
Comparative shell and soft-body morphology of Turritriton? sp. 1 from Kwajalein Atoll, Marshall Islands (images courtesy of S. Johnson, n.d.; Kwajalein Underwater Website). Left: Three views of two individual shells (top: 23.0 mm; bottom: 24.0 mm), showing coarsely sculptured axial ribs, dark reddish-brown coloration, and prominent peripheral nodules. Right: Live-animal images of the corresponding specimens, highlighting darker pigmentation and distinct mantle patterning in comparison to Turritriton labiosus sensu stricto. This morphotype is rare in the Marshall Islands and has been found under rocks and among Halimeda algae at depths of 6–16 m. Its overall morphology is notably similar to that of the Tenerife specimen.

The close morphological alignment of the Tenerife specimen (Fig. 1) with these Indo-Pacific forms (Figs. 2–5) raises compelling questions regarding the species’ biogeographic origin in the eastern Atlantic.

**Figure 5.**
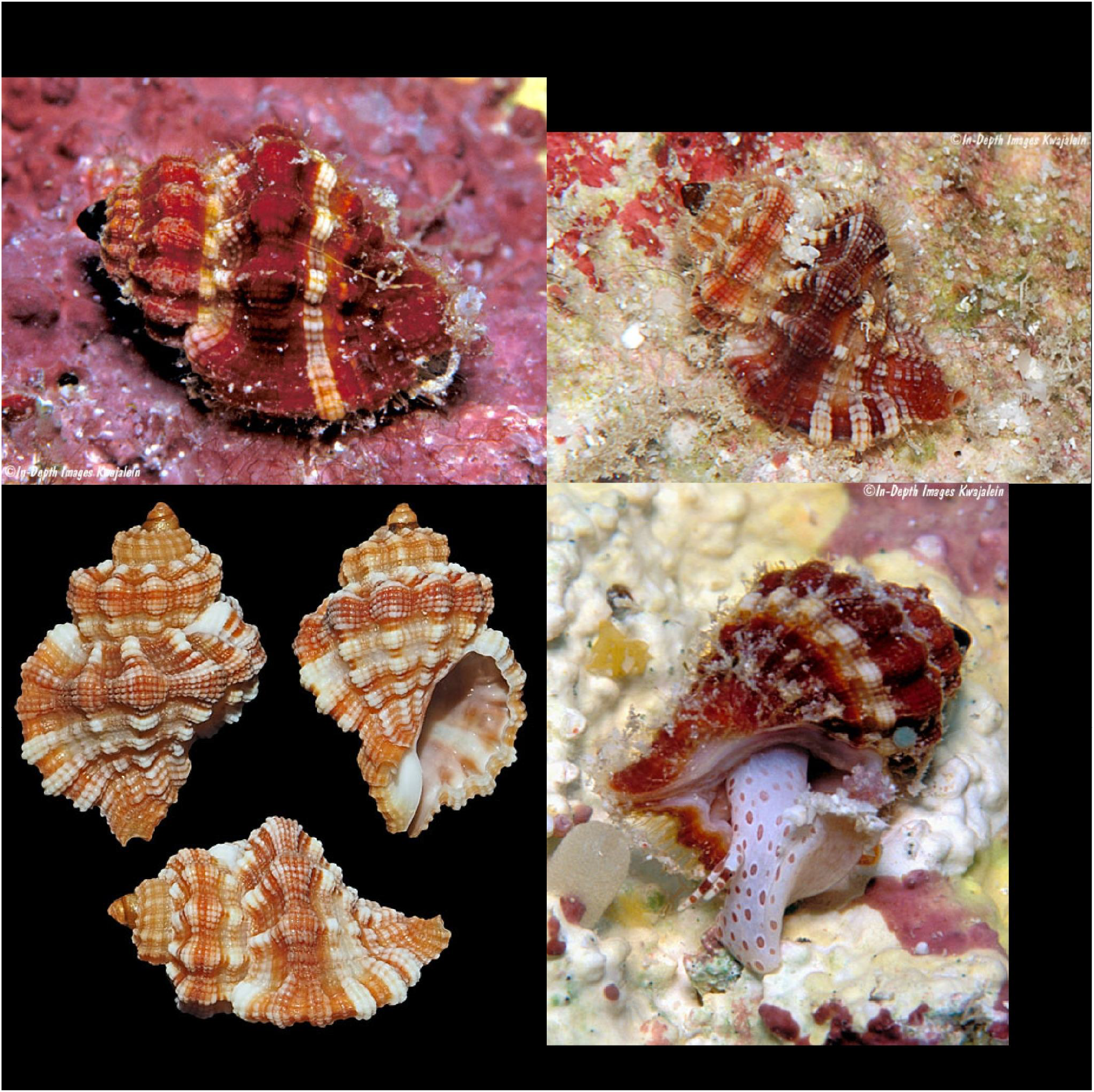
Shell and live-animal views of *Turritriton labiosus* (Wood, 1828) sensu stricto from Kwajalein Atoll, Marshall Islands (images courtesy of S. Johnson, n.d.; Kwajalein Underwater website). Top and right: In situ photographs of living individuals with consistent shell morphology—orange-red pigmentation, narrow axial ribs, and fine spiral cords. Mantle coloration is pale with diffuse reddish to purple spotting. Bottom left: A representative shell (17.2 mm) of the typical Indo-Pacific form, exhibiting finely developed axial sculpture, compact fusiform shape, and white subsutural bands. This morph is moderately uncommon in the Marshalls and typically inhabits Halimeda-rich microhabitats at 6–16 m depth.

Natural long-distance dispersal via a planktotrophic larval stage is biologically plausible, particularly considering prevailing equatorial and subtropical currents and the extended pelagic larval durations documented for ranellid gastropods. Alternatively, anthropogenic vectors—such as ballast water discharge or rafting on marine debris—may have facilitated its introduction. The existence of both historical and previously undocumented shells—including the first modern published record from Lanzarote (Vicente, 2007)—further supports the hypothesis of a small, possibly cryptic population rather than a single, transient occurrence.

Johnson (2020) highlighted notable intraspecific variation among T. labiosus specimens from the Marshall Islands, including differences in axial sculpture and soft-part pigmentation. The Tenerife shell (Fig. 1) closely resembles the “Sp. 2” morphotype from Kwajalein Atoll (Fig. 3), which Johnson regarded as potentially distinct based on consistent differences in both shell architecture and live-animal coloration. This raises the possibility that the Atlantic specimen belongs to a previously unrecognized lineage within a cryptic Indo-Pacific–Atlantic species complex.

To resolve the taxonomic and biogeographic uncertainties presented here, integrative approaches—including anatomical studies and molecular phylogenetics—are required. Broader sampling across the Macaronesian archipelagos, combined with comparative genetic analysis and DNA barcoding of Indo-Pacific material, will be essential to determine whether the Canary Islands population represents a recent colonization event, a disjunct relict lineage, or a previously overlooked and undescribed Atlantic species.

This new record extends the known range of Turritriton labiosus, contributes a valuable data point to the biogeography of Cymatiidae, and emphasizes the importance of ongoing biodiversity monitoring in oceanic island systems, where Indo-Pacific taxa may occasionally establish viable extra-limital populations. The morphological data presented here contribute to the assessment of intraspecific variation and support the hypothesis of a geographically structured species complex. These findings underscore the value of integrative taxonomy in clarifying species boundaries and distributional patterns within Ranellidae sensu lato.

## Supporting information

- A Darwin Core `occurrence.csv` file - `meta.xml` mapping file - `eml.xml` metadata file

## Acknowledgments

I am grateful to Bob Abela and Scott Johnson for generously providing photographic material and for permitting its use in this study. Their documentation of Indo-Pacific Turritriton specimens significantly contributed to the comparative analysis presented here. I also wish to express my sincere appreciation to Luigi Bozetti, Werner Massier, Danilo Gubbioli, Frank Swinnen, Andrea Nappo, David Monsecour, Lacopo Briano, Armando Verdasca, Ricardo Vega, Clemente Rebora and Christian Girard for their enduring support of regional malacological research and for their valuable contributions to fieldwork, specimen documentation, and collaborative exchange. This study was supported by Coralliophila.com, Tenerife.

## References

Abela, B. (2018, October 20). Turritriton labiosus (W. Wood, 1828), Apra Harbor, Guam [Photograph]. In Guam Ranellidae – Guam Gastropods. Accessed June 12, 2025, from: https://www.bobabela.com/Guam-Mollusks/Guam-Gastropods/Guam-Ranellidae/i-8ZMHBSb

Abela, B. (2018, October 21). Turritriton labiosus from Apra Harbor, Guam [Photograph]. International Seashell Collectors Club. Accessed June 12, 2025, from: https://www.facebook.com/photo.php?fbid=10218072425707601

Beu, A. G. (2010). Neogene tonnoidean gastropods of tropical and South America: Contributions to the revision of Indo-Pacific Ranellidae. Bulletins of American Paleontology, 377–378: 1–550.

Biota. (2025). Turritriton labiosus (Wood, 1828).

Biodiversidad de Canarias. Accessed June 11, 2025, from: https://www.biodiversidadcanarias.es/biota/especie/E02629

Bommerer, M. (2025). Turritriton labiosus observation. iNaturalist. Accessed June 12, 2025, from: https://www.inaturalist.org/observations/283931969

Conchology.be (2015). Turritriton labiosus specimen from the Philippines. Shell presented with dorsal and ventral views; collection data unspecified. Accessed June 12, 2025, from: https://conchology.be/?t=68&u=619311&g=08faab7acb8780b3075dd0d9cbad7e44&q=c872300133357711a4128c8832803c3d

García-Talavera, F. (1983). Estudio faunístico y biogeográfico de los moluscos marinos del Archipiélago Canario. Doctoral thesis, Universidad de La Laguna, Tenerife, 648 pp.

García-Talavera, F. (1987). Malacofauna del Archipiélago Canario: Moluscos marinos. Consejería de Educación y Cultura del Gobierno de Canarias, 381 pp., 17 pls.

iNaturalist (2025). Turritriton labiosus observations in the western Atlantic. Accessed June 12, 2025, from: https://www.inaturalist.org/observations?nelat=34.697705809881015&nelng=-54.894029104107375&subview=map&swlat=7.788693534376527&swlng=-87.32566972910737&taxon_id=488923

Johnson, S. (2020, July 24). Two morphotypes of Turritriton labiosus from Kwajalein Atoll [Discussion with images]. Living Sea Shells, Facebook. Accessed June 12, 2025, from: https://www.facebook.com/groups/138530896852326/permalink/595784141126997/

Johnson, S. (2020). Turritriton labiosus observation. iNaturalist. Accessed June 12, 2025, from: https://www.inaturalist.org/observations/109315029

Johnson, S. (n.d.). Turritriton labiosus (as Cymatium labiosum). Kwajalein Underwater. Accessed June 12, 2025, from: https://www.underwaterkwaj.com/shell/triton/Cymatium-labiosum.html

Johnson, S. (n.d.). Turritriton? sp. 1. Kwajalein Underwater. Accessed June 12, 2025, from: https://www.underwaterkwaj.com/shell/triton/Cymatium-sp1.html

Kapella (2023). Turritriton labiosus (Wood, 1828). Image from Tenerife. Image archived in MolluscaBase. Accessed June 12, 2025 from: https://www.marinespecies.org/aphia.php?p=image&pic=154915&tid=476594

MolluscaBase eds. (2024). Turritriton labiosus (W. Wood, 1828). Accessed June 22, 2025, through: World Register of Marine Species at: https://www.molluscabase.org/aphia.php?p=taxdetails&id=476594

Nordsieck, F. & García-Talavera, F. (1979). Moluscos marinos de Canarias y Madeira (Gastropoda). Cabildo Insular de Tenerife, Tenerife, 208 pp.

RedPROMAR (n.d.). Red de Observadores del Medio Marino en Canarias. Gobierno de Canarias. Accessed June 12, 2025, from: https://www.redpromar.org/

Vicente, J.L. (2007). The family RANELLIDAE Gray, 1854 (GASTROPODA: TONNOIDEA) in the Canary Islands, with special emphasis on Lanzarote. Visaya, 2(2), 17–38. https://www.academia.edu/40466664/The_Family_RANELLIDAE_Gray_1854_GASTROPODA_TONNOIDEA_in_the_Canary_Islands_with_special_emphasis_on_Lanzarote

von Cosel, R. (1982). Beiträge zur Kenntnis der marinen Molluskenfauna von Ghana. Archiv für Molluskenkunde, 112(1–3): 43–100.

WoRMS Editorial Board. (2024). Pagurus anachoretus Risso, 1827. Accessed June 12, 2025, through: World Register of Marine Species at https://www.marinespecies.org/aphia.php?p=taxdetails&id=107231

